# An algorithm for the identification of indicator taxonomic units and their use in analyses of ecosystem state

**DOI:** 10.1101/2022.05.16.492087

**Authors:** Hernán de la Vega, Liliana Falco, Leonardo Saravia, Rosana Sandler, Andrés Duhour, Víctor N Velazco, Carlos Coviella

## Abstract

Biological community structure can be used as an ecological state descriptor, and the sensitivity of some taxonomic groups or biological entities to environmental conditions allows for their use as ecological state indicators. This work describes a mathematical methodology developed for the identification of such taxonomic units when comparing environments or ecosystems under different anthropic impacts. Based on this methodology, a freely downloadable R package for easy use was developed, (Ecoindicators, <u>DOI: 10.5281/zenodo.5772829</u>). Based solely on presence or absence information, the method identifies indicator taxonomic units for each environment, estimates the belonging of any additional samples to a given environment, approximates the ecological niche of any taxonomic unit based on two or more selected environmental factors, and returns the frequency of any taxonomic unit in a selected combination of environmental factors. By using the approximation to the ecological niche of species present, given a new sample, the physicochemical parameters can be estimated by the species present in the sample. These analyses can be performed simultaneously for two or more taxonomic units. This paper provides a mathematical description of how the mathematical method was developed. One of the advantages of this method, and the referred R-package is that it can be used for any ecosystem for which there is a suitable biological dataset associated with environmental factors. In addition, both this mathematical procedure and the package referred to, can be tailored by other researchers to fit their own needs. We also expect other developers to further improve it.

## 1 Introduction

The development of biological indices of environmental status as a tool to assess anthropic impact is increasingly used in many systems (Melo-Merino, 2020). These biological status indices are well developed for aquatic environments but their development for terrestrial environments is still incipient (Guerra et al., 2021; Huera-Lucero et al., 2020; Nunes et al., 2020; Rocha et al., 2020). The European Water Framework Directive, for example, required that all surface waters in Europe have biological indices of water quality by 2015 (EEA 2018 European Parliament, 2000). However, the development of reliable ecological quality indices requires not only the identification of those biological units considered to be indicators, but also the development of objective methodologies for the construction of such indices. A current characteristic of the development of these indices is the general lack of standardized tools and methodologies for the objective selection of variables and for their construction (Velásquez et al., 2007). The purpose of this work is to advance in the design of such unbiased tools and the methodologies that can be used for the construction of ecological system state indices. Thus, we designed an algorithm to classify the most relevant characteristics of ecosystems and estimate the values of parameters considered of interest, using the presence and absence of certain taxonomic units. Such units considered here as the biological entities of different taxonomic hierarchy used in this work, from samples of the same system. The work started from a database of soil samples that contain measurements of physical and chemical parameters as well as the presence and absence in each sample of different taxonomic units as defined above. The samples were obtained over two years of sampling in sites of different intensity of anthropic use of the same soil. The identification of the different biological units that make up the edaphic biota, as well as their interactions and dynamics, are difficult to assess due largely to the methods necessary for their extraction and the small size of the individuals that compose it. However, the information gradually collected over decades, is reaching the point where it is becoming possible to focus the work on the development of comparative studies on the structure and functioning of the edaphic biota. These studies will then make possible the analysis of the stability of the interaction networks for evaluating the state of different ecological systems (Fortin et al., 2021; Lau et al., 2017) or of the same system under different intensities of anthropic impact (Potapov et al., 2019). As a first step, certain taxonomic units were selected, called here “indicators” that were then used (observing their presence or absence) to estimate from which environment a soil sample came. As a second step, the presence or absence of such units was used to estimate values of some physical and chemical parameters of interest. To carry out this task in an automated way with different databases, an algorithm was developed that allows to complete all these stages in a single step. The first problem addressed was to determine, when receiving new soil samples, to which environment they correspond. The focus of interest in this part, was to make this classification taking into account those units present or absent, regardless of the values of the chemical and physical parameters of the samples. With that aim, it was first sought to distinguish “indicator units” that, through their presence or absence, increase or decrease the probability that a sample belongs to a specific environment. With this information, and observing only those presences or absences in new soil samples, the algorithm estimates which environment they come from and assigns a probability to that estimation. In the following section this procedure is detailed and in the final section a test is carried out with a database corresponding to a soil of the rolling pampas (Buenos Aires, Argentina). A second objective consisted in relating the presence of taxonomic units (indicator units) in the samples with the levels of certain physical and chemical parameters of interest. This problem was tackled by describing the intersection of the ecological niches *sensu* Hutchinson (1957) of the groups present with respect to those parameters.

To this end, it was necessary to choose a simple calculation to obtain an approximation of the niches. A “grid” of the ranges of the physical and chemical parameters of the database was built and then a “convex capsule” from a representative part of the existing cloud of biological data was adjusted. This whole process is described in the last section. It is also intended for the entire procedure to be written in a free language known to researchers in the area, with the intention that it can be tested by other professionals and improved by other developers.

### Methodology: construction of the algorithm step by step

The database has the structure (Figure 1) in which the columns are completed with measurements of physical and chemical parameters and of the gross abundances for each taxonomic unit present in each of the samples obtained from a same type of soil with different intensities of anthropic impact.

**Figure 1.**
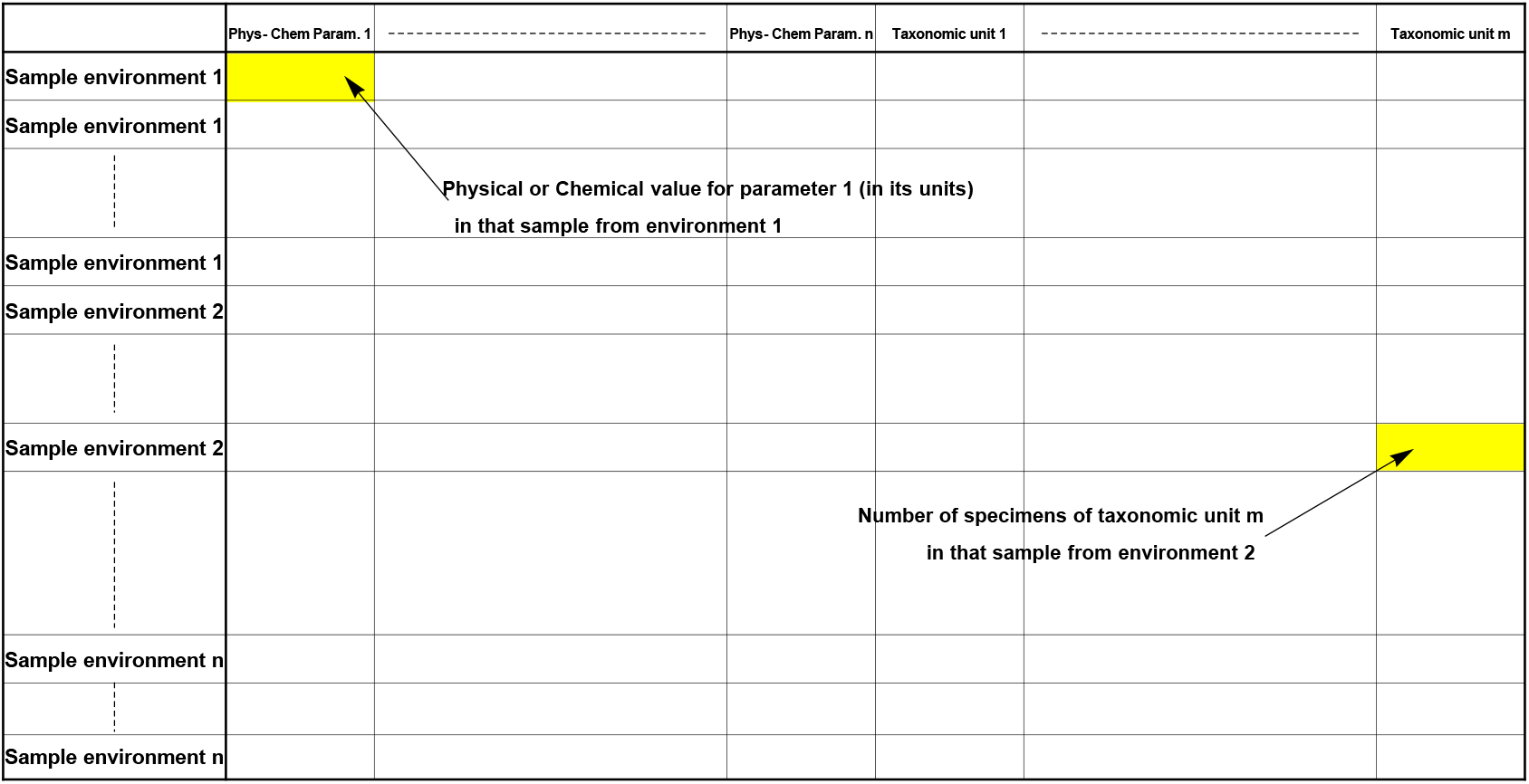
Database structure. The samples come from the same soil subjected to three different intensities of anthropic use (environment). Each line contains the physicochemical data and the taxonomic units found in each sample

## 2 Procedure for estimating a sample belonging to a given environment

From the samples obtained from an experimental design and reaching the laboratory, the assignment probability that relates a sample to a particular environment is calculated. This step, to calculate the probability of assignment, begins by considering the presence / absence (Figure 2) of the taxonomic units present in the sample.

**Figure 2.**
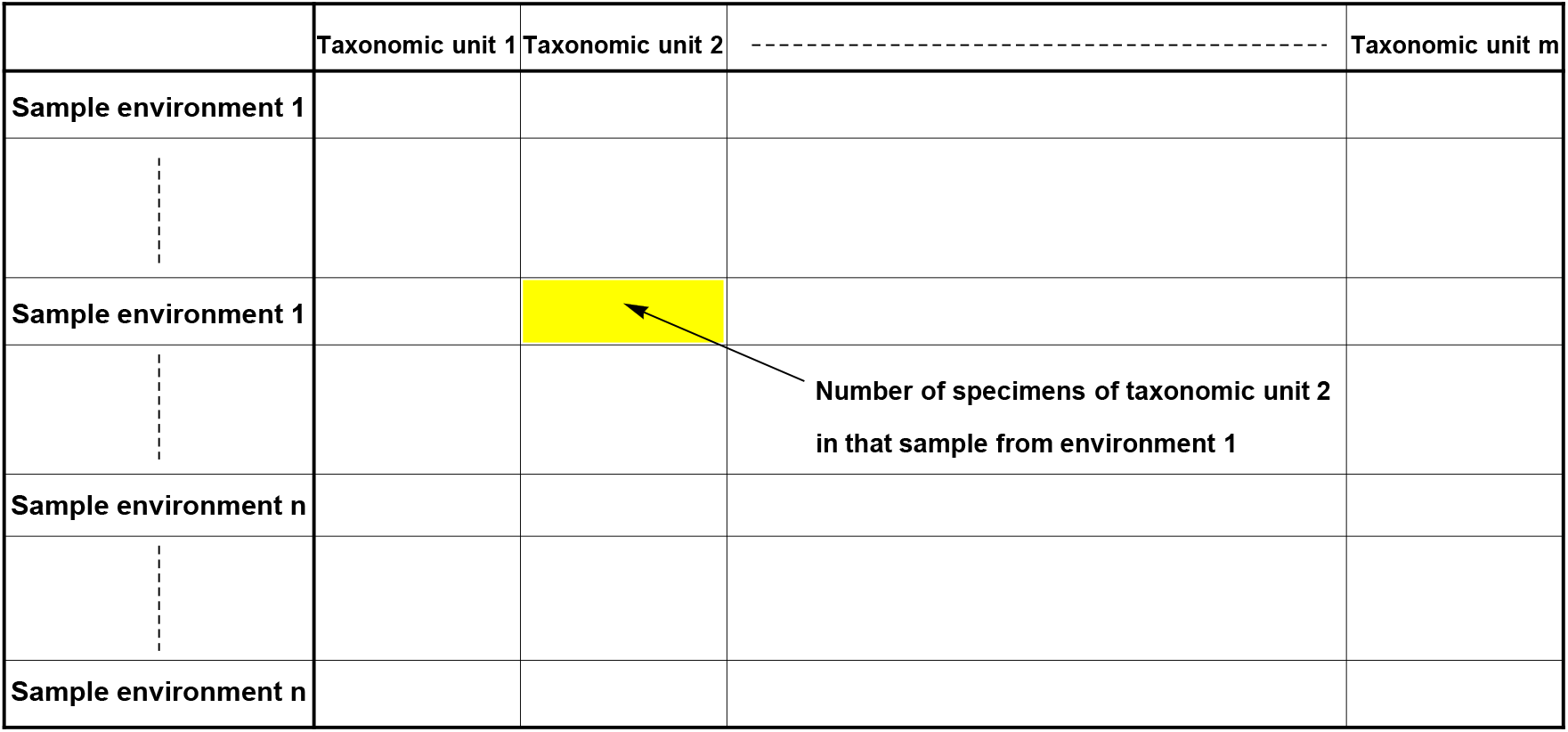
Trimmed table, containing only information about the taxonomic units present or absent. Each row is a sample with the values corresponding to the number of individuals of each taxonomic unit found in that sample.

**Figure 3.**
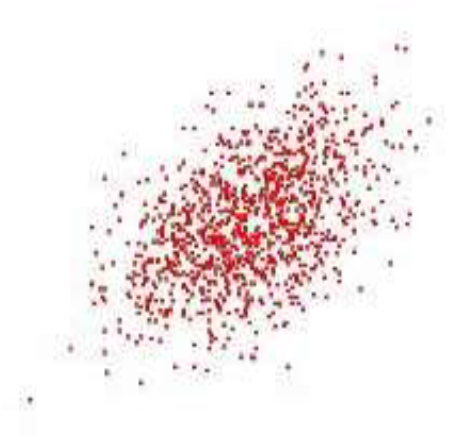
The entire point cloud according to 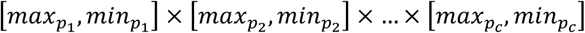

As indicated above, the algorithm considers the presence or absence of each taxonomic unit, therefore, the columns of the taxonomic units obtained from the database are transformed into a matrix of zeros and ones. In this first version, the number of samples from each environment is required to be the same. So, the first matrix obtained is:

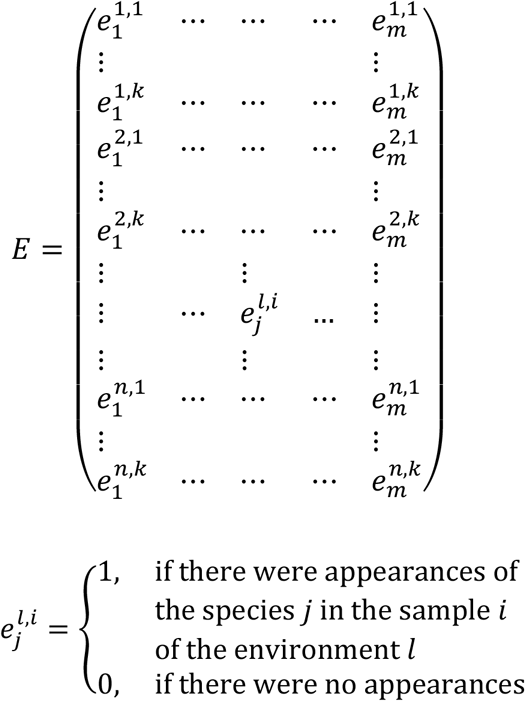

Where the term 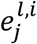indicates if there were appearances of the species *j* in the sample *i* of the environment *l*. More precisely:

### 2.1 Selection of indicator unit

The idea that is being tested here is to measure the difference between the expected and observed values. As the number of samples *k* corresponding to each of the environments *n* is the same, if the appearance of each taxonomic unit *j* were independent of the environments, it would be expected that the proportion of appearances of each taxonomic unit in each environment was uniform. To specify the latter, let’s take a taxonomic unit *j* and add its occurrences in the environment *l*, then add all its occurrences in the database, take the quotient between the two and call that number 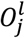that is:

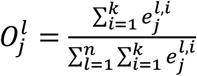 were 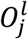gives the proportion of appearances observed, with respect to the total of appearances of the unit *j*, in the environment *l*.

If the unit *j* were independent of the different environments, would be expected then that it your occurrence 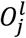 be approximately 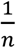 the same for each environment *l* = 1, …, *n* this is 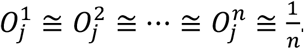.

We call 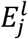this expected value and we have that 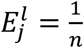for *l* = 1 … *n* and *j* = 1 … *m*. We test the hypothesis 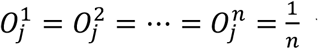with a standard Hypothesis Test using the Chi-square distribution *χ*^2^ by:

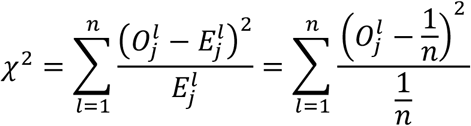

If the value exceeds the threshold given by *α* = 0.05, the hypothesis is rejected and it is considered that the occurrences of that taxonomic unit vary between environments.

Units whose distribution is not uniform with respect to the environments are called Indicator Units, and are those in which their occurrences are not independent of the environments being considered. A number *r* of indicator units is thus obtained 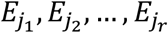.It is through their appearances or absences the procedure seeks to determine the belonging of a new sample to a certain environment.

To visualize this, the algorithm generates a graph with the distributions of each unit in each environment and indicates the number of total occurrences of each one in the database (Figure 6, section 5.1). This last datum is considered in the calculation above so as not to use units that appeared only a few times to be of significance in the analysis.

### 2.2 Environment estimation

Once the Indicator Units have been obtained, we seek to determine which particular environment it belongs to. Specifically, the negation of the Null Hypothesis test (with *alpha* = 0.05): “the observed proportion of that unit in that environment is 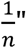 is used to construct the *D* matrix based on the differences between the expected value 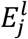and the observed occurrence rate 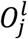, so that each 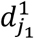is equal to 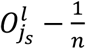

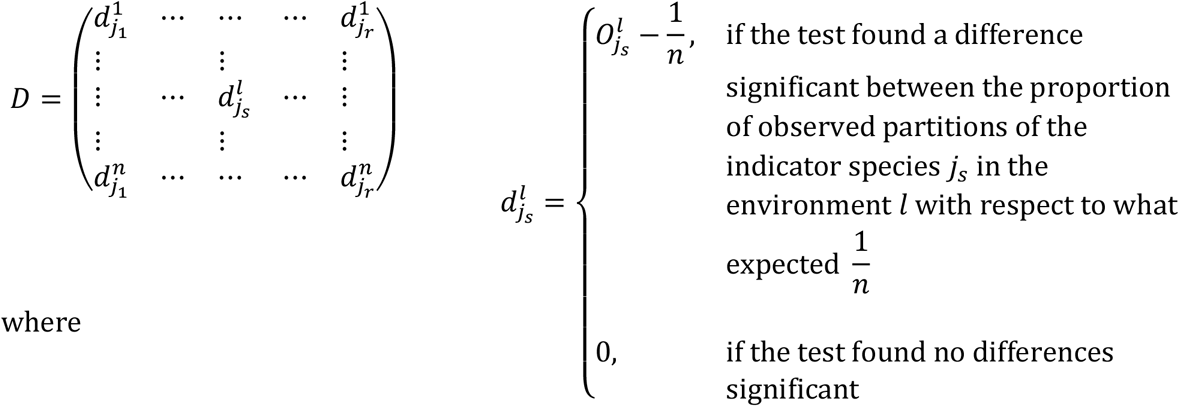

The rationale is that these coefficients 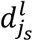, before the appearance of a unit in a sample *j*_*s*_, add or subtract probabilities (or neither of those two things) this new sample belongs to a certain environment. For instance, suppose there are three different environments and the algorithm has selected two indicator units. Furthermore, assume that the proportions of occurrences of each indicator unit in each environment with respect to its total occurrences (the values 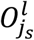),

|  | Indicator unit 1 | Indicator unit 2 |
| --- | --- | --- |
| Environment 1 | 0.35 | 0.10 |
| Environment 2 | 0.60 | 0.62 |
| Environment 3 | 0.05 | 0.28 |

Suppose additionally that the tests carried out on each unit in each environment to see if the observed proportions deviate from those expected (which in this case would be 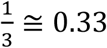 gives a negative result for Indicator Unit 1 in Environment 1 and for Indicator Unit 2 in Environment 3, then the array of values 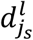would be (rounding the values):

|  | Indicator unit 1 | Indicator unit 2 |
| --- | --- | --- |
| Environment 1 | 0 | -0.23 |
| Environment 2 | 0.27 | 0.29 |
| Environment 3 | -0.28 | 0 |

where that 0 of Indicator Unit 1 in Environment 1 indicates that the test did not find a significant difference between what was observed and what was expected, and that 0.27 of Indicator Unit 1 in Environment 2 indicates that the test did find a significant difference. The algorithm then takes that difference 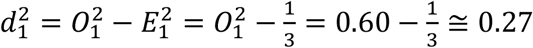, and repeats the procedure with the other values of the matrix. A concrete example of the construction of this matrix is given in Figure 7 of section 5.1.

As stated before, the idea is that these numbers add or subtract probabilities that a sample belongs to a certain environment. For example, if Indicator Unit 1 appears in a new sample, Environment 2 will add 0.27 points to the probability of that sample belonging to that environment while Environment 3 would subtract 0.28 points.

The way the procedure uses that information is as follows. Suppose that a number *q* of new samples are received, all from the same environment, an environment that the procedure seeks to identify. To continue with the previous example (with three Environments and two Indicating Units) suppose that we receive four samples in which the following quantities of each Indicating Unit are recorded in each Environment:

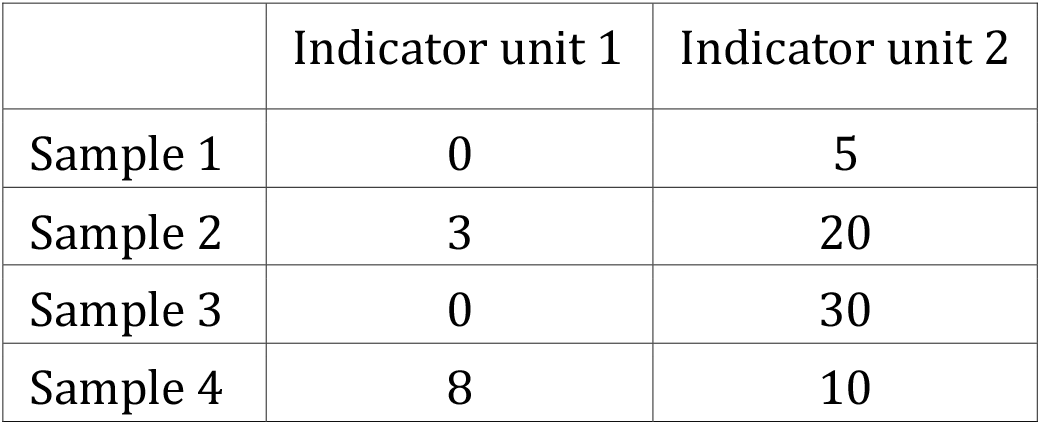

The matrix is again translated so that it only contains presences and absences:

|  | Indicator unit 1 | Indicator unit 2 |
| --- | --- | --- |
| Sample 1 | 0 | 1 |
| Sample 2 | 1 | 1 |
| Sample 3 | 0 | 1 |
| Sample 4 | 1 | 1 |

Preserving the subscripts that have been used for the indicator units, this matrix would then have the form:

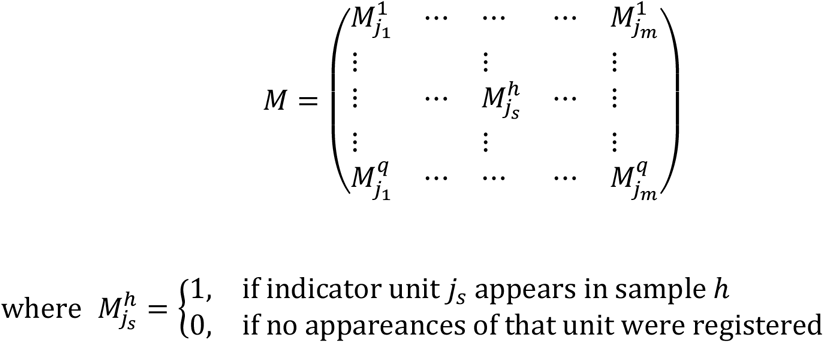

A vector of occurrences of the samples is then built that contains, in each coordinate, several of the samples received where there were occurrences of each unit:

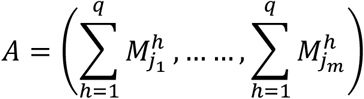

In the example provided it would be *A* = (0 + 1 + 0 + 1,1 + 1 + 1 + 1) = (2,4)

Then the values of the matrix *D* should be added, (which increase or decrease the chances of belonging to a certain environment) according to the number of samples in which there were appearances of each Unit. It is then calculated *R* = *A. D*^*t*^ that it is the product between the matrix (the vector) *A* and the transpose of the matrix *D*. In the example:

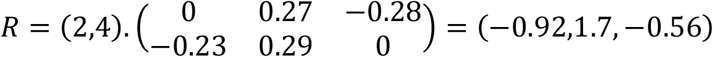

Each coordinate of the vector *R* corresponds to one of the environments, in the example:

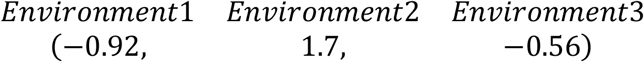

The largest of these coordinates indicates the environment the procedure is looking for. That is, the procedure considers that the set of samples corresponds to the environment whose coordinate has the highest value in the vector *R*. In the example it corresponds to Environment 2.

### 2.3 Estimations validation

Even before receiving new samples of the system, it is of great interest to test the operation of the algorithm. One strategy for this is to take the original database and take some random samples from it from the same environment, as if they were new samples, run the algorithm and verify if it is correct in the prediction, errs in the prediction or it is not able to give an answer regarding which environment the samples belong to. Let‘s go back to the matrix *E*. The rows in this matrix contain all the samples in the database and in each of the lines there are zeros or ones according to whether or not there are occurrences of each unit in the base. Here, the algorithm separates this matrix into three sub-matrices containing each of the samples from a single environment, as:

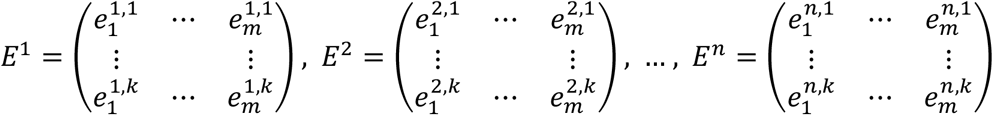

The algorithm now randomly chooses one of these matrices and a random sample from that matrix, runs the process as if it were a new sample and determines if it was correct in the prediction, erred in the prediction or could not give a prediction. It repeats this process n times and calculates the percentage of hits, misses, and no prediction results. Subsequently, it repeats the process described above, but this time taking two samples from each environment (instead of one). Then it takes three samples from each environment and so on up to thirty samples from each environment or the maximum possible if the database does not contain that many samples per environment. The algorithm then returns three graphs (hits, misses and no prediction) displaying the previous calculations, which gives an idea of the accuracy of the predictions. In the example shown in the section 5.1 the resulting graphs are shown for the example given here (Figure 8).

## 3 Estimation of physical and chemical parameters from the units present

When trying to link biological, physical and chemical data, several problems immediately appear, some of them quite obvious: The number of variables is usually very large and the necessary computing power exceeds the capacity of the available resources. The choice of the way forward becomes difficult if one wants to simplify the problem.

### 3.1 Niche approximation

For its practical application, the interest is usually focused on relating only a limited number of physical and chemical parameters with only some of the units. Thus, we select a quantity “*c*” of physical and chemical parameters that here are called “*p*_1_, *p*_2_, …, *p*_*c*_” and a taxonomic unit “*e*”. If in a sample of the database there is an occurrence of the unit “*e*”, a vector (*v*_1_, *v*_2_, …, *v*_*c*_) can be built with the values registered in that sample of the parameters “*p*_1_, *p*_2_, …, *p*_*c*_“. Performing this task for all samples in which there is an occurrence of the unit taxonomic “*e*”, we obtain a “cloud” of points in space *R*^*c*^.

In the cases *c* = 1,2,3 that point cloud can be graphed.

Now let’s take the ranges in which each parameter of interest moves, in this way we obtain for each parameter “*p*_*j*_” a value 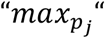 and a value 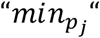 with which a “hypercube” is constructed that contains the entire point cloud that represents the presence of the taxonomic unit “*e*”.

The hypercube containing the total point cloud for the taxonomic unit “*e*” can be segmented by choosing a number “*d*” that will divide each interval 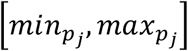obtaining smaller hypercubes called *d*_*c*_ in which *c* is the number of selected physical-chemical parameters and *d* is the number of times each range of the physical-chemical parameters will be divided. The choice of these values will be subject to the computing power of the computers and can be selected based on the bibliography and the researcher’s criteria.

In the case *c* = 3, of the three-dimensional case shown, there are many algorithms capable of constructing the “convex capsule” of the point cloud as a way to approximate the niche of the species with respect to the three particular parameters chosen.

The point cloud can be observed within different squares (figure 4) of resolution “*d*” representing the hypercubes “*d*_*c*_”. The number of occurrences of the taxonomic unit “*e*” in each hypercube “*d*_*c*_” is recorded to obtain a “fit” value from the point cloud.

**Figure 4.**
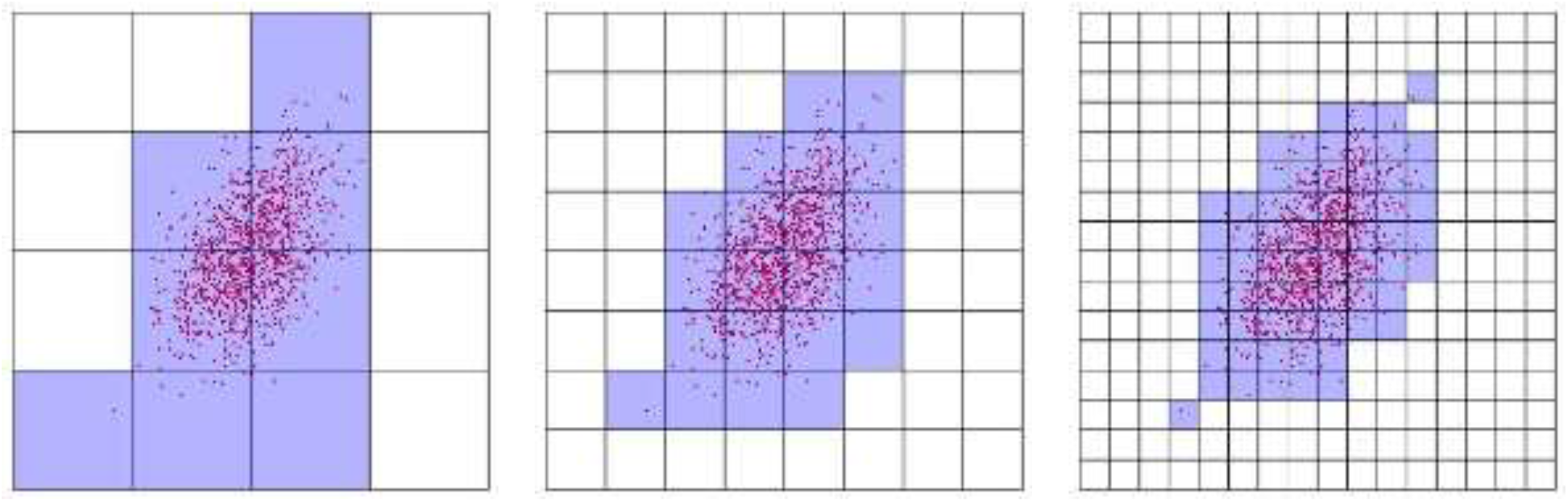
Different resolutions of the point cloud according to the size of the cells or grid resolution.

### 3.2 Parameter estimation

Point clouds for the occurrence of two taxonomic units taking into account the same physicochemical parameters, co-occur in a few hypercubes (figure 5). Then the intervals can be limited under observation of the physicochemical parameters.

**Figure 5.**
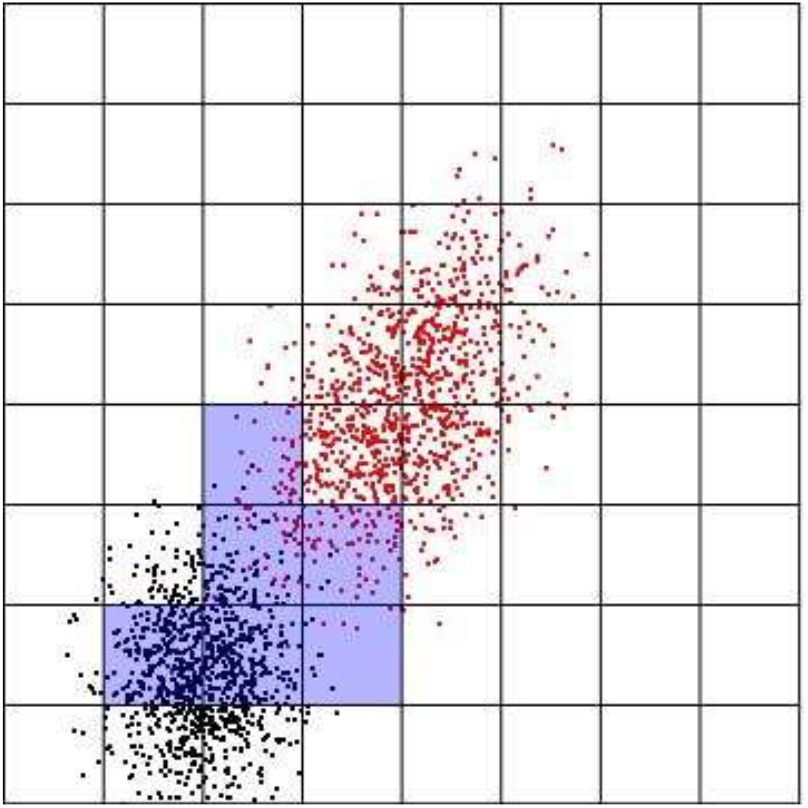
Superposition of the cloud of points corresponding to two taxonomic units and a grid that represents hypercubes “*d*_*c*_”. The shaded area corresponds to the hypercubes in which the joint appearance of both taxonomic units occurs.

The algorithm distinguishes the hypercubes where there were appearances of the taxonomic units by collecting and ordering the information. Hypercubes are labeled by *d*_*c*_

Here we tried two different approaches:

In the first one, for each taxonomic unit, the occurrence number for each hypercube is obtained.

An “occurrence” matrix is created that shows the number of occurrences of each unit in each cube. In this matrix, the rows *m* are the database samples and the columns are the hypercubes *d*_*c*_, so each value *a*_*i,j*_ indicates the number of samples in which the unit *i* appears in the *j* cube. *m*

This matrix is tedious to calculate and requires time. For this reason, the algorithm exports the matrix obtained as a csv file and later reads the values directly from that file.

The algorithm, on the one hand, returns which taxonomic units appear in a sample and in which hypercube “*d*_*c*_” they are found.

On the other hand, given a number of taxonomic units then it returns the samples, from the database, in which all those taxonomic units appear at the same time and in which cubes they appear.

Also, given a number of taxonomic units, it returns how many times all those units appear together in each cube.

Each of the described steps are used to create a function that returns the hypercubes where the taxonomic units occur and the probability of being in a certain cube (knowing that all those units appeared in that sample).

As a second approach, two vectors *m* are created (one for each species) of *d*^*c*^ coordinates (one for each cube). Each one of these vectors, contains either a 1 if that particular species appears at least once in a given cube, or a 0 if it never appeared in that cube. These vectors are easier to calculate than the matrix “appearance” described above, because it contains less information. These approach saves processing time and computer resources, but some other calculations cannot be performed this way.

If we then choose a certain number of species and want to visualize in which cubes they appear together, we only need to obtain the product, coordinate by coordinate, of the vectors of each species, and look in which coordinate each species appear.

## 4 Indicator units within the grid

Indicator units are characterized by appearing more in certain environments than in others (Dufrene et al.,. Proceeding as in the previous section, those physicochemical parameters of interest are chosen and the corresponding grid is made, with which more information can be obtained on how the difference between the expected and observed proportion of the units in each environment occurs. If we observe one indicator unit within the grid, the difference between the expected and the observed proportions in each cube can be calculated, which allows to visualize cubes (or “zones”, sets of cubes) where the difference is greater. To do this:

A function is created that indicates the number of times and the percentage in which each cube appears in each environment.

The number of times and the percentage in which each cube appears in the entire database is calculated (without discriminating between environments).

With the above data, a matrix is built that has in each row the number of times each cube appears in each environment and in total.

Given a unit, the number and percentage of occurrences in each cube discriminated by environment are recorded (the number and percentage of occurrences in each cube having already been calculated without discriminating by environment).

With the previous data, a matrix is constructed that in each row shows the number of occurrences of the unit in each cube, discriminated by environment and in total.

The next step is to build a matrix called” Projections” that shows an estimate of how many times the unit should appear in each cube in each environment, assuming that its appearances were independent of the environment. Specifically, if we call:

- *c*_*i,j*_ to the number of times the cube appears *i* in the environment *j*
- *a*_*i*_ to the number of appearances of the unit in the cube *i* in total (without discriminating by environment)
- *c*_*i*_ to the number of times the cube appears *i* in total (without discriminating by environment)

then the Projection matrix has in place the (*i,j*) value 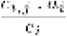 as an estimate of the number of times the unit should appear in the cube *i* in the environment *j*

The difference between the last two matrices is then calculated, and shows the difference between the observed and the expected appearances of the unit in question in each cube and in each environment.

This difference is also calculated as a percentage.

These differences are displayed using histograms. This visualization becomes more relevant as long as the number of cubes is not too large.

## 5 A concrete example of how the algorithm works

### 5.1 Estimation of parameters from the units present

This section shows a concrete example of the use of the described process and the calculations and results obtained for that case. The database used in this example corresponds to a soil from the Pampean plain (Buenos Aires, Argentina). Each sample in this database collects measurements of fifteen physical and chemical parameters and the presence or absence of forty-three taxonomic units. The database has 216 samples in total corresponding to three different environments (72 samples from each environment). Environment 1 corresponds to a naturalized grassland (NG), Environment 2 to a grazing field that passed onto agriculture two years before the start of the samplings (CG), and Environment 3 is an environment of continuous intensive agriculture for at least 40 years (AG). The procedure begins with the selection of the indicator units, for them the matrices described in the section2 are calculated. Here the graph is shown where the difference between the observed and expected occurrences expressed as percentages is observed. In this case, as there are three environments, the value of the expected proportions (if the units were independent of them) is 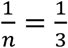 and it is represented by a horizontal line.

Figure 6. All taxonomic units labeled with numbers are shown. Total number of occurrences of the tagged taxonomic units. The vertical axis represents the rate of occurrence of a taxonomic unit, the limit 0.33 is the expected value 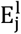. Vertical dotted lines are those indicator species. Environments are represented by colors. Environment 1 = black Environment 2 = red Environment 3 = green. The units selected as indicators are Rhodacaroidea; Parasitoidea; Veigaioidea; Euphthiracaroidea; *Microscolex_dubius; Eukerria_stagnalis*.

**Figure 6.**
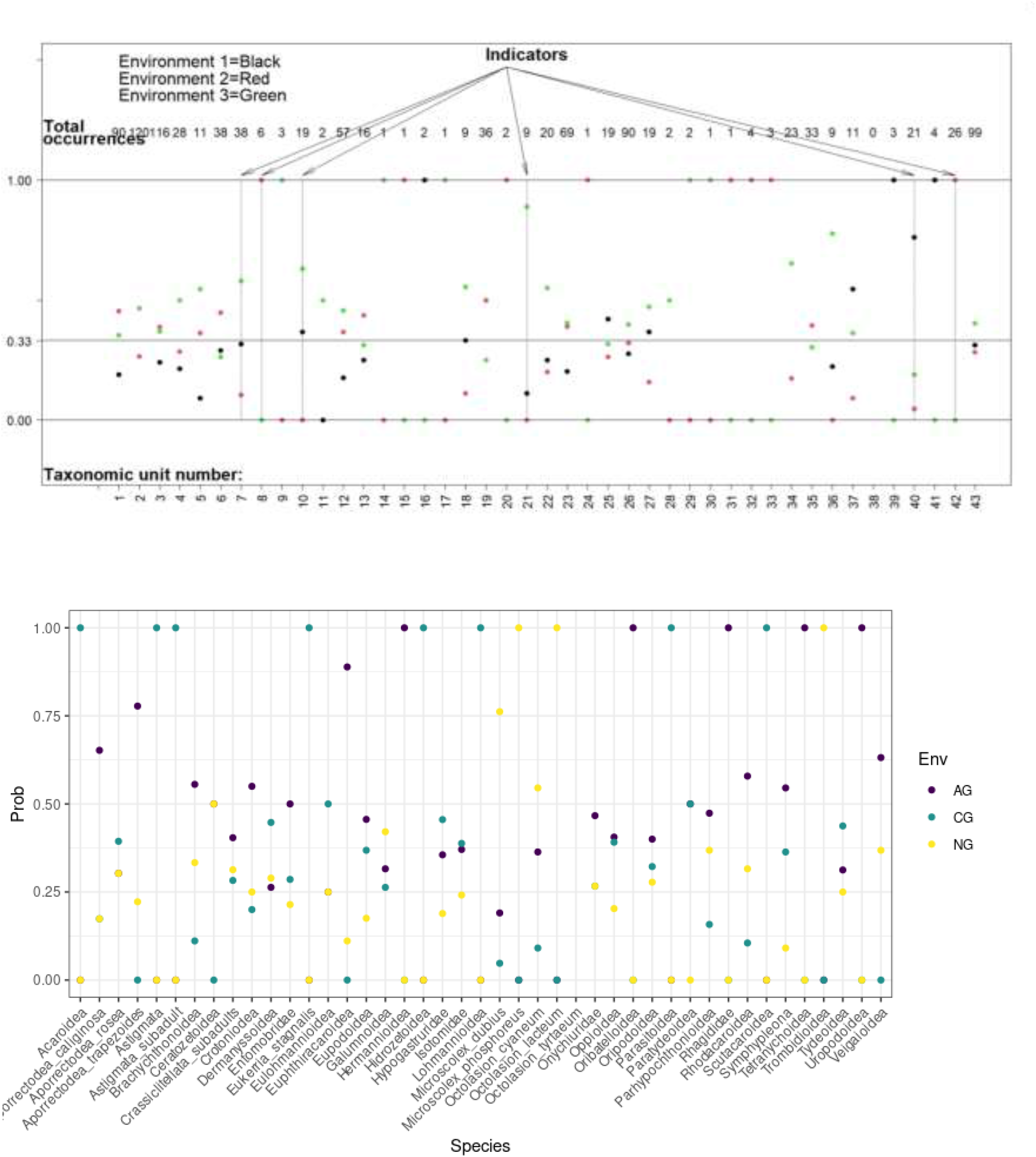
shows the observed proportions in each environment, and they are represented by color dots.

The procedure goes as in section 2.2 producing a coefficient matrix *D* (Figure 7):

**Figure 7.**
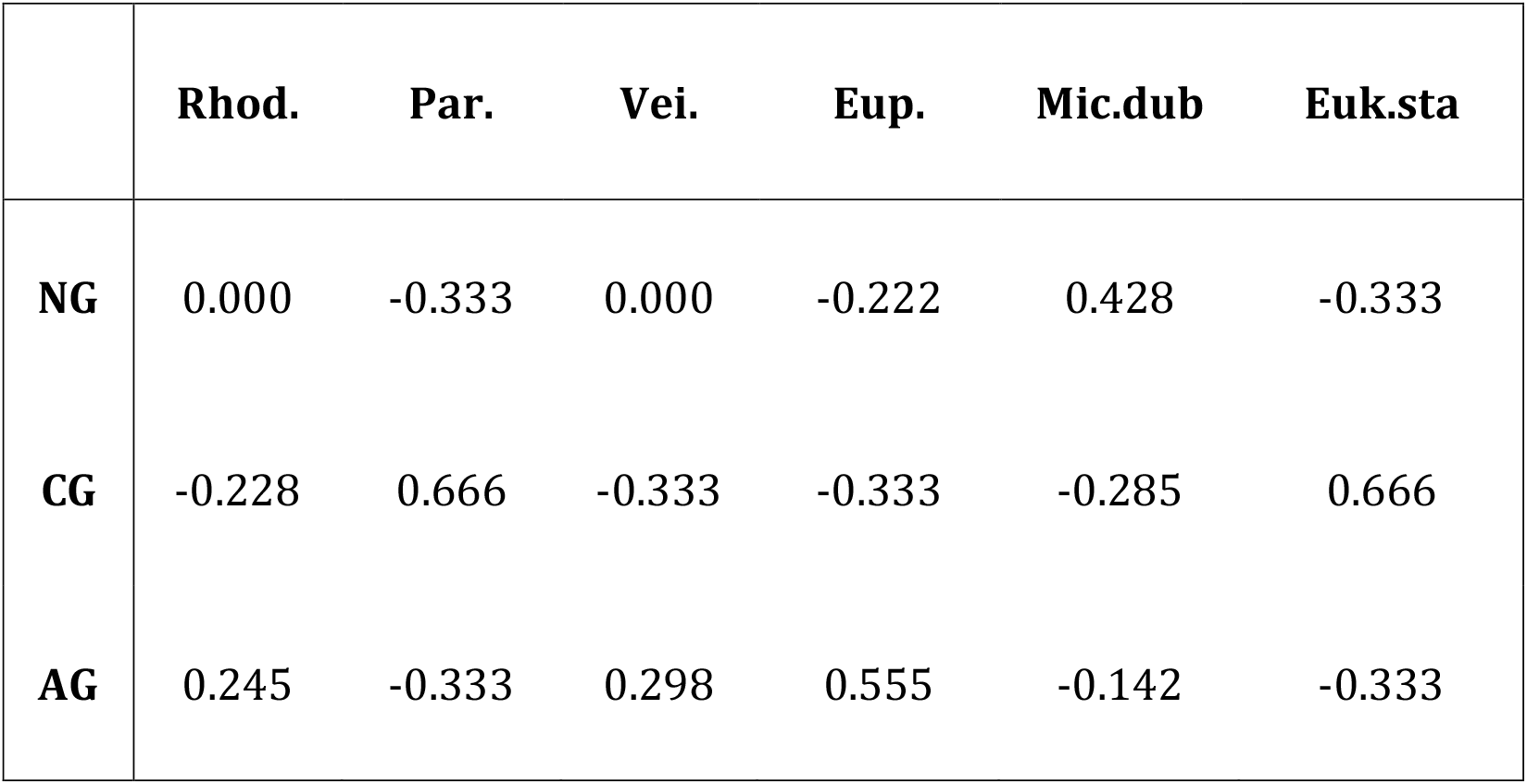
Coefficient matrix *D* of the units selected as indicators.

Now suppose that samples of the same type of soil were received but about their environments (or management) we do not know, and this process is used to determine which environment/management they belong to. As described in section 2.2, we take the coordinates corresponding to the indicator units and they are replaced by 0 if there were no occurrences of the units in that sample and 1 if there were. Then:

|  | **Rhod.** | **Par.** | **Vei.** | **Eup.** | **Mic.dub** | **Euk.sta** |
| --- | --- | --- | --- | --- | --- | --- |
| **Sample 1** | 0 | 0 | 0 | 0 | 1 | 0 |
| **Sample 2** | 1 | 0 | 1 | 0 | 0 | 0 |
| **Sample 3** | 0 | 0 | 1 | 0 | 1 | 0 |

For instance, sample numbers 27, 35, and 36 of the database give these results and all three belong to the same NG environment. The values of the samples are added:

|  | **Rhod.** | **Par.** | **Vei.** | **Eup.** | **Mic.dub** | **Euk.sta** |
| --- | --- | --- | --- | --- | --- | --- |
| **Sum** | 1 | 0 | 2 | 0 | 2 | 0 |

The product between this last vector is then carried out with the transpose of the matrix *D*:

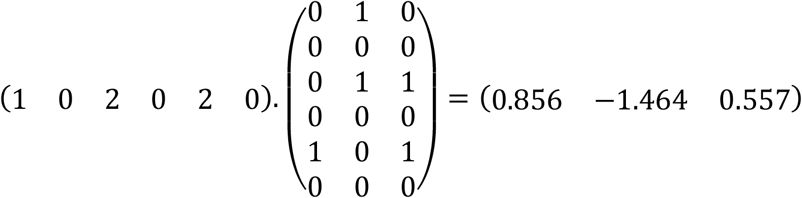

That is:

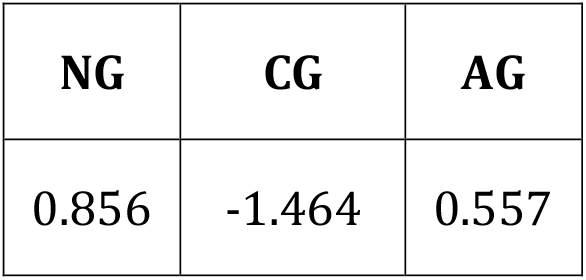

| **NG** | **CG** | **AG** |
| --- | --- | --- |
| 0.856 | -1.464 | 0.557 |

As the largest of the numbers corresponds to the NG environment, it is concluded that the samples come from that environment.

In this case, the prediction coincides with the actual origin of the samples. As described in section 2.3, this process was carried out several times with one sample, with two samples, with three samples, and so on. The percentages of hits, misses, and times in which the algorithm cannot decide which environment the set of received samples belongs to are then calculated. The percentages are shown in Figure 8.

**Figure 8.**
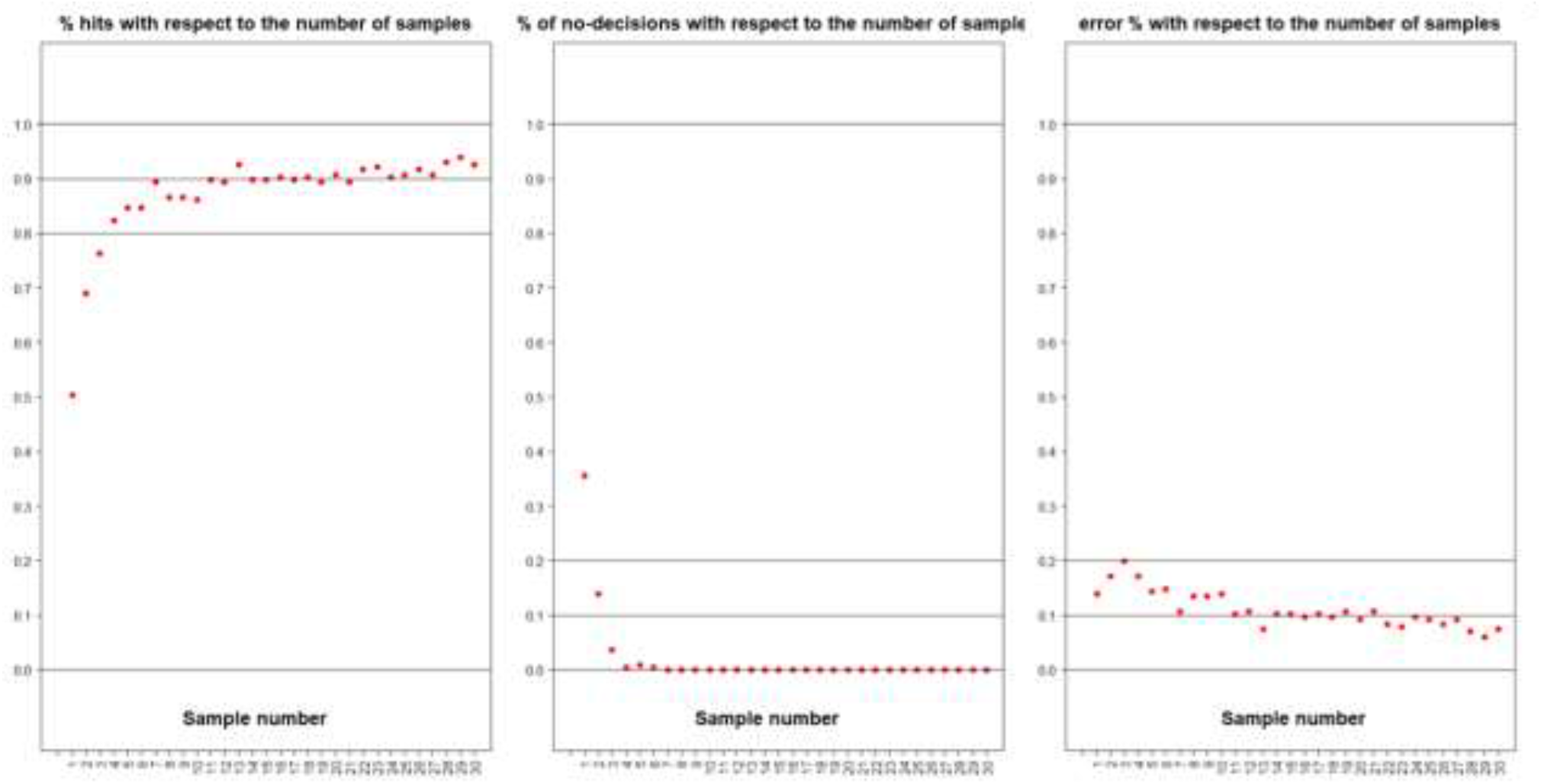
Percentages of hits, no-decisions, and misses calculated.

### Estimation of parameters from the units present

In this example, the physical-chemical parameters that have been chosen (Sandler, 2019) were P, OM and N. It can be observed that the minimum and maximum values recorded for P are 0.00 and 75.78; those corresponding to OM are 1.51 y 9.2, and those of N are 0.14 and 0.51.

Each of these ranges is divided into 3 parts (*d* = 3) and a “grid” formed by 27 cubes is obtained as shown in figure 9.

**Figure 9.**
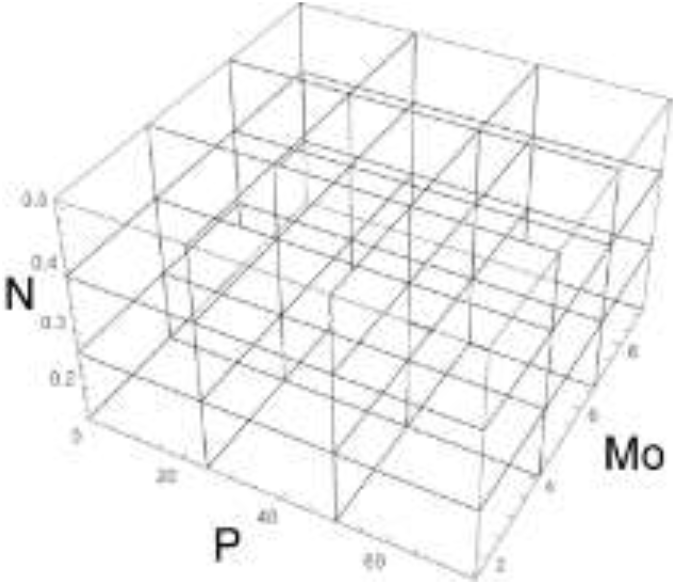
Cube built with Nitrogen (N), Phosphorous (P), and Organic matter (OM) environmental factors.

**Figure 10.**
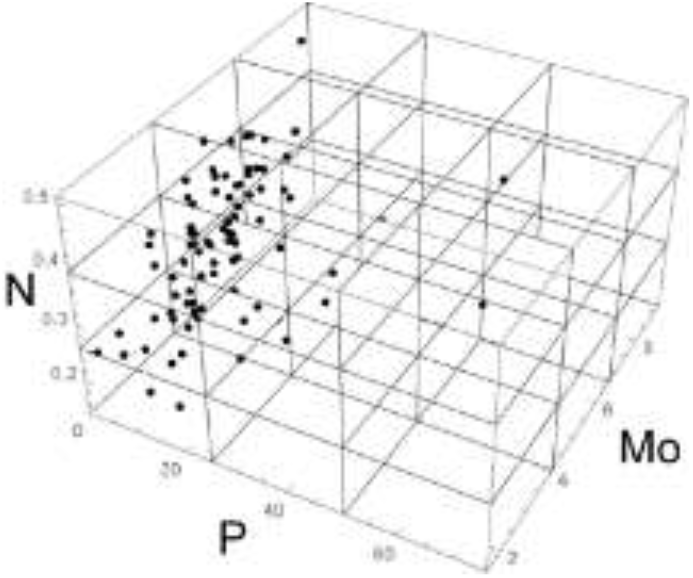
Cube built with N, P, and OM, showing the presence of the Onychiuridae taxon in a grid of the environmental variables N, P and OM.

From the observation of the appearances of each unit within the grid, the procedure goes as described in section 3.2.

For example, the simultaneous appearance of the units “Onychiuridae”, “Isotomidae”, “Eupodoidea”, and “*Aporrectodea rosea*” is detected only in samples that appear in the cube delimited by 0.00 ≤ *P* < 25.26, 4.08 ≤ *Mo* < 6.66 and 0.26 ≤ *N* < 0.39 (Figure 11)

**Figure 11.**
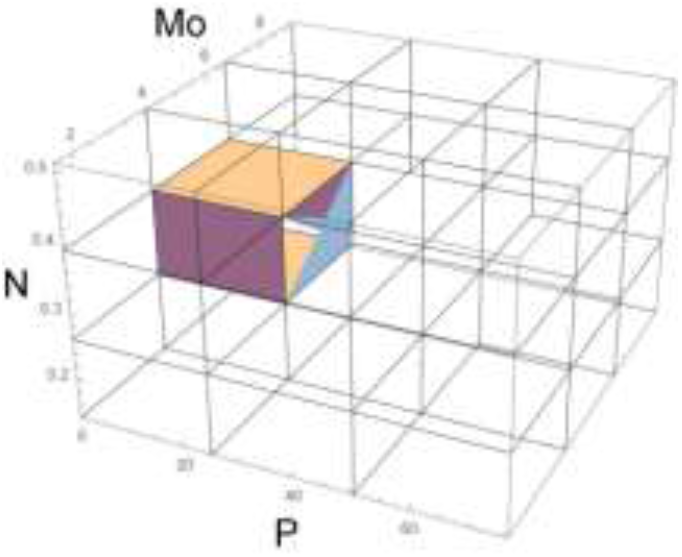
Simultaneous occurrence of taxonomic units “Hypogastruridae”, “Crotoniodea”, and “Crassiclitellata subadults”.

The simultaneous appearance of the units “Hypogastruridae”, “Crotoniodea”, and “Crassiclitellata subadults” is only detected in samples that appear in the cube delimited by 0.00 ≤ *P* < 25.26, 4.08 ≤ *Mo* < 6.66 and 0.14 ≤ *N* < 0.26 and additionally in the cube delimited by 0.00 ≤ *P* < 25.26, 0.00 ≤ *P* < 25.26 and 0.26 ≤ *N* < 0.39. Joining both cubes it can then be determined a zone of simultaneous appearances delimited by 0.00 ≤ *P* < 25.26, 4.08 ≤ *Mo* < 6.66 and 0.14 ≤ *N* < 0.39.

### Indicator units within the grid

Let’s now take the parameters P, OM and N as in the previous subsection and build the same division into cubes. Let us also take the indicator unit “Rhodacaroidea” and compare its distribution in each cube in each environment with the expected distribution if the appearances of the unit were independent of the different environments.

The three graphs that are in the upper part of Figure 13 show how many times that unit appears in each cube and each environment (cubes numbered here from 1 to 27). The three graphs in the lower part show how many times it should appear under the hypothesis of independence of environments.

**Figure 12.**
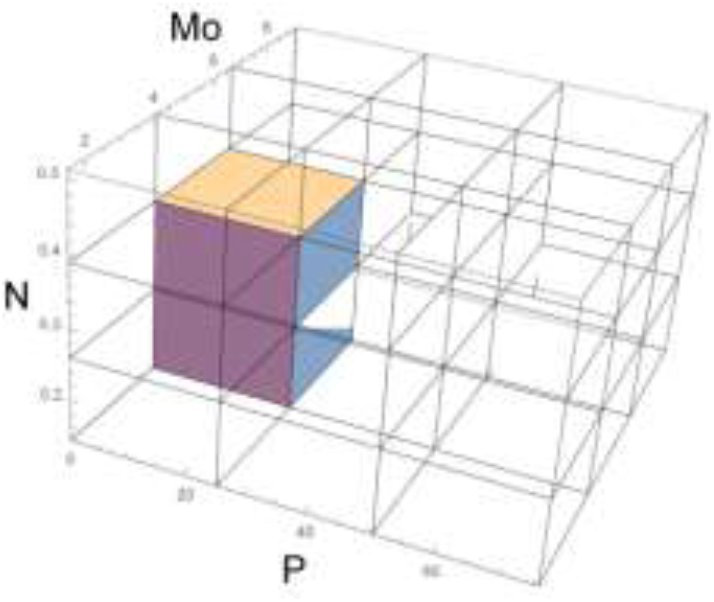
Simultaneous occurrence of taxonomic units “Onychiuridae”, “Isotomidae”, “Eupodoidea”, and “*Aporrectodea rosea*”.

**Figure 13.**
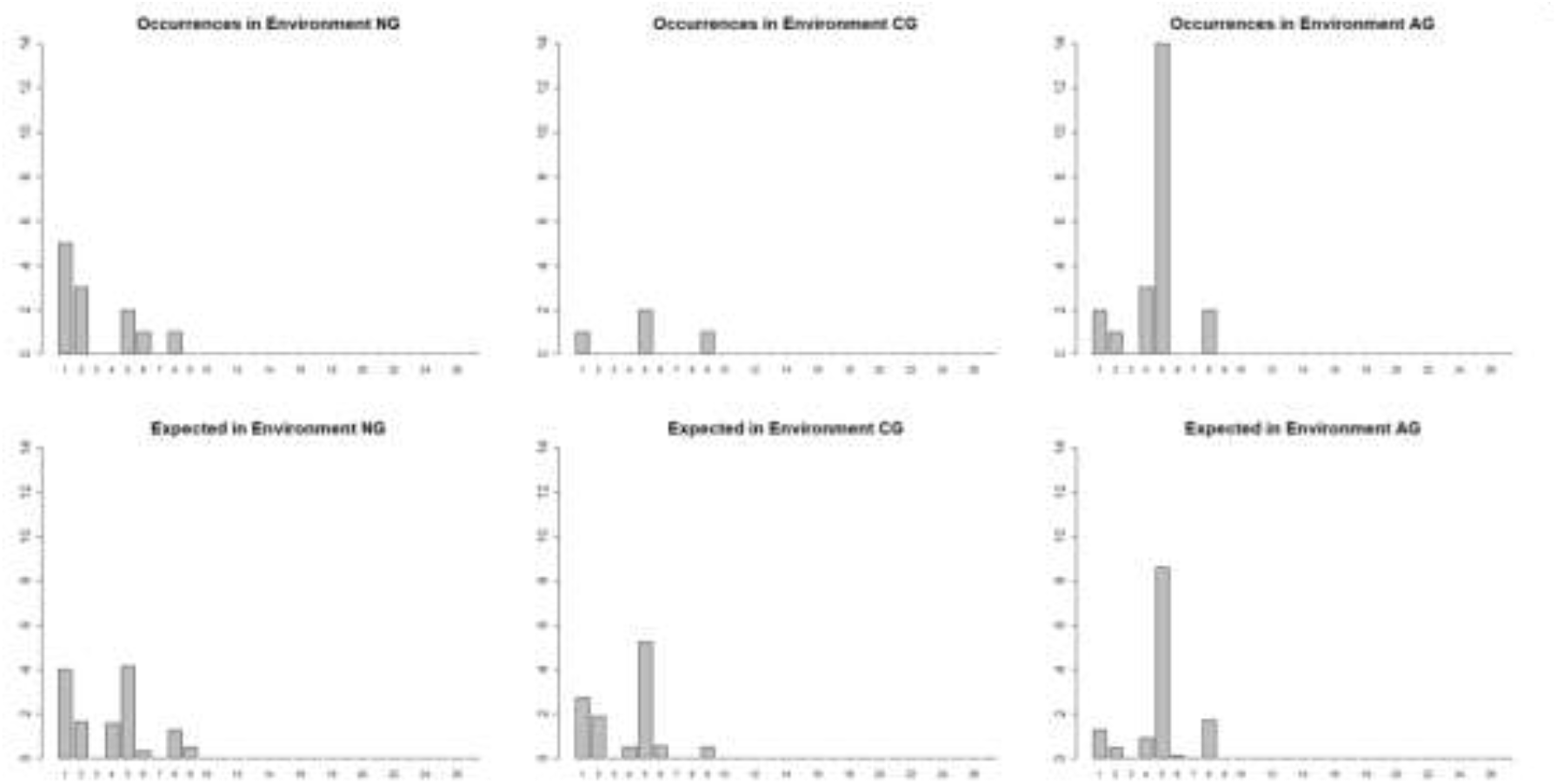
Observed (upper panels) and expected (lower panels) occurrence of unit Rhodacaroidea in the three environments being compared. Natural grassland (NG), Cattle grazing (CG), and Agriculture (AG).

It can be seen in Figure 6 that the unit “Rhodacaroidea” (with the number 7) appears more frequently than expected (which in the example is 30 percent) in environment 3 (environment AG), less frequently than expected in environment 2 (CG) and with the frequency expected in environment 1 (NG). These differences between what is observed and what is expected can be found in some aspects of Figure 13. It can be seen, for example, that in the cubes that appear with the numbers 1, 4, and 5, there is a marked difference upwards (between expected and observed) in the AG environment and a marked difference downwards in the CG environment.

## Conclusions

Although this method was originally developed using a soil biota database (Sandler, 2019), it will work just the same in any other environment or ecosystem for which there is a suitable database of biological entities, associated with an environmental dataset. The algorithms developed and put together in this new method not only allow for the identification of indicator taxonomic units (depending on the taxonomic resolution available to the user), but also to approximate their ecological niches and, given a new sample, to estimate the physicochemical parameters of the site according of the species present in that sample.

One of the main advantages of this method is that it can be used for any ecological system for which there is a suitable biological dataset associated to environmental factors. Another useful feature, is that it requires only presence/absence data. Researchers that also have density data, can modify and improve the method by tailoring it to their datasets. Thus, we feel that this contribution will be of interest for researchers developing indicators of ecosystem state. Moreover, the entire procedure was then converted to “Ecoindicators” (de la Vega et al., 2019), an R package that performs all the required tasks. The package (<u>DOI: 10.5281/zenodo.5772829</u>) is free to use and to be improved by any researcher, with proper citation.

## Bibliography

de la Vega H., Falco L., Saravia L., Sandler R., Duhour A., Coviella, Carlos. (2019). Ecoindicators. DOI: 10.5281/zenodo.5772829.

Dufrêne, M. & Legendre, P. Species Assemblages And Indicator Species:the Need For A Flexible Asymmetrical Approach Ecological Monographs,1997, 67, 345–366.

EEA. European Environmental Agency Report 7. 2018. European waters. Assessment of status and pressures /2018. ISBN 978-92-9213-947-6 ISSN 1977-8449 doi:10.2800/303664

Fortin M-J, Dale MRT, Brimacombe C. 2021 Network ecology in dynamic landscapes. Proc. R. Soc. B 288: 20201889. https://doi.org/10.1098/rspb.2020.1889

Guerra, C. A., Bardgett, R. D., Caon, L., Crowther, T. W., Delgado-Baquerizo, M., Montanarella, L. & Eisenhauer, N. (2021). Tracking, targeting, and conserving soil biodiversity. Science, 371(6526), 239–241.

Huera-Lucero, T., Labrador-Moreno, J., Blanco-Salas, J., Ruiz-Téllez, T. (2020). A Framework to Incorporate Biological Soil Quality Indicators into Assessing the Sustainability of Territories in the Ecuadorian Amazon. Sustainability 12, 3007; doi:10.3390/su12073007

Hutchinson, G. E. 1957. Concluding remarks. Cold Spring Harbor Symposium in Quantitative Biology. 22:415–427.

Lau, M. K., S. R. Borrett, B. Baiser, N. J. Gotelli, and A. M. Ellison. (2017). Ecological network metrics: opportunities for synthesis. Ecosphere 8(8):e01900. 10.1002/ecs2.1900

The Scientific World Journal 2021, 12 pages, https://doi.org/10.1155/2021/9970957.

Nunes, M., Karlen, D., Veum, K., Moorman, T., Cambardella, C. (2020). Biological soil health indicators respond to tillage intensity: A US meta-analysis. Geoderma 369, 114335 https://doi.org/10.1016/j.geoderma.2020.114335.

Potapov AM, Tiunov AV, Scheu S. (2019). Uncovering trophic positions and food resources of soil animals using bulk natural stable isotope composition: Stable isotopes in soil food web studies. Biol. Rev.; 94, 37–59. https://doi.org/10.1111/brv.12434

Rocha, L., Hegoburu, C., Torremorell, A., Feijoó, C., Navarro, E., Fernández, H. (2020). Use of ecosystem health indicators for assessing anthropogenic impacts on freshwaters in Argentina: a review Environmental Monitoring Assess 192:611 https://doi.org/10.1007/s10661-020-08559-w

Sandler, Rosana V. (2019). Indicadores de sustentabilidad del suelo basados en la estructura y funcionamiento de la fauna edáfica. Ph.D. dissertation. Universidad Nacional de General Sarmiento. Argentina.

Velásquez, E., Lavelle P., Andrade M. (2007). GIQS: a multifunctional indicator of soil quality. Soil Biology & Biochemistry. 39: 3066–3080. DOI:10.1016/JSOILBIO.2007.06.013

